# An expanded genetic toolbox to accelerate the creation of *Acholeplasma laidlawii* driven by synthetic genomes

**DOI:** 10.1101/2022.09.21.508766

**Authors:** Daniel P. Nucifora, Nidhi D. Mehta, Daniel J. Giguere, Bogumil J. Karas

**Affiliations:** Department of Biochemistry, Schulich School of Medicine and Dentistry, The University of Western Ontario, London, ON N6A 5C1, Canada

**Keywords:** *Acholeplasma laidlawii*, electroporation, replicative plasmids, synthetic cell, genome transplantation, strain evolution

## Abstract

Assembling synthetic bacterial genomes in yeast and genome transplantation has enabled an unmatched level of bacterial strain engineering, giving rise to cells with minimal and chemically synthetic genomes. However, this technology is currently limited to members of the Spiroplasma phylogenetic group, mostly *Mycoplasmas*, within the *Mollicute* class. Here, we propose new genetic tools for developing these technologies for *Acholeplasma laidlawii*, which is phylogenetically distant from *Mycoplasmas* and, unlike most *Mollicutes*, uses a standard genetic code. We first investigated a donor-recipient relationship between two *A. laidlawii* strains through whole-genome sequencing. We then created multi-host shuttle plasmids and used them to optimize an electroporation protocol. We also demonstrated the use of evolution to create superior strains for DNA uptake via electroporation. For genome transplantation, we selected *A. laidlawii* 8195 as the recipient strain and created a PG-8A donor strain by inserting a Tn5 transposon carrying a tetracycline resistance gene. The tools presented here will improve *Acholeplasma* research and accelerate the effort toward creating *A. laidlawii* strains driven by synthetic genomes.

**Graphical Abstract:** 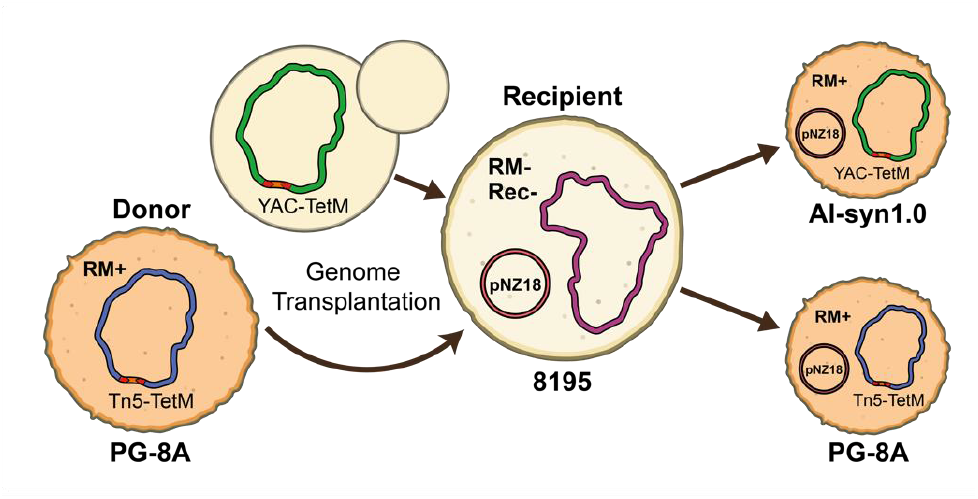

## INTRODUCTION

Synthetic biology is a rapidly growing field that aims to modify or create organisms for use in such avenues as industry, biotechnology, and agriculture. Recently, the development of two breakthrough technologies has taken the first step towards harnessing the full potential of bacterial species through the generation of synthetic/highly engineered strains. The first technology is the cloning of whole bacterial genomes in a eukaryotic host such as *Saccharomyces cerevisiae*^1^. Once cloned or assembled in *S. cerevisiae*, the yeast genetic toolbox offers species-independent genome editing that often exceeds what is possible in the original bacterial system^2,3^. The second technology is a method of whole-genome transformation termed ‘genome transplantation’. This method involves the use of polyethylene glycol (PEG)-mediated transformation to deliver a ‘donor’ genome (i.e., one that has been isolated from another bacterium or one that has been cloned or assembled in yeast) into a closely related recipient bacterium^4^. The resulting transformed cell will assume genotypic and phenotypic traits conferred by the donor genome. Combining these technologies, highly engineered bacterial strains, including synthetic, minimized, and attenuated organisms, have been created^5–7^. Although extremely powerful, genome transplantation is currently limited to bacteria within the class *Mollicutes*, particularly those in the genus *Mycoplasma*^8,9^.

*Phytoplasmas* are plant pathogenic *Mollicutes* that can cause symptoms such as yellowing, stunting, and phloem necrosis, which can result in decreased crop yields^10^. Laboratory research of *Phytoplasmas* is heavily restricted to date: only recently have some strains been successfully cultured in axenic media^11,12^. One way to overcome this problem is to create hybrid organisms that carry *Phytoplasma* genomes but also can be propagated in laboratory conditions. Since *Phytoplasmas* are likely too distantly related to *Mycoplasmas* for genome transplantation to be successful^8^, we propose *Acholeplasma laidlawii*, a close relative to *Phytoplasmas*^13^, as a platform for creating hybrid synthetic organisms. Additionally, *Acholeplasma* and *Phytoplasma* species are among the only *Mollicutes* to use a standard genetic code, meaning *Phytoplasma* genes can be expressed in *A. laidlawii* without the need for recoding (most *Mollicutes* use the UGA codon to encode tryptophan). Furthermore, unlike some *Mollicutes, A. laidlawii* does not require expensive serums and has fewer biosafety restrictions. Multiple *A. laidlawii* strains have been studied since the late 1900s, and perhaps the most extensively studied is strain PG-8A. This strain has been used as a representative of the *Acholeplasma* genus, having a sequenced genome, mapped proteome, and characterized promoter structure^14,15^. The genome of strain PG-8A has also been cloned in *S. cerevisiae*^16^. However, despite the extensive research involving this strain, virtually no genetic tools exist for manipulation *in vivo*, and even reports of transformation are very limited^17^.

Other strains of *A. laidlawii* have been more amenable to genetic manipulation. For instance, strain 8195 is a restriction-deficient derivative of *A. laidlawii* JA1, which has a history of use for the propagation and study of Mollicute-infecting viruses^18,19^. Strain 8195 has more genetic tools available, including transformation, replicative and transposon-bearing plasmids, and heterologous gene expression^20–22^, and is therefore a good candidate for a recipient cell that would allow for rebooting synthetic hybrid strains. Despite this past work with strain 8195, many of these tools, and the knowledge for using them in *A. laidlawii*, have seemingly been lost from the scientific community.

Here, we show the development of a new OriC plasmid for *A. laidlawii* 8195, an improved electroporation protocol, and the creation of PG-8A donor strains with Tn5-transposase insertion. Our genetic toolbox presented here is the first step towards enabling genome transplantation between *A. laidlawii* strains PG-8A and 8195, which will ultimately lead to the creation of synthetic hybrid *Acholeplasma* strains.

## RESULTS AND DISCUSSION

To enable the creation of *A. laidlawii* strains driven by synthetic genomes, an efficient way to transfer DNA between various strains needs to be developed. Specifically, whole-genome transfer from bacteria to host organisms, as well as genome transplantation from donor to recipient bacterial strains must be established. The genetic tools required for achieving these tasks are listed in Figure 1.

**Figure 1.**
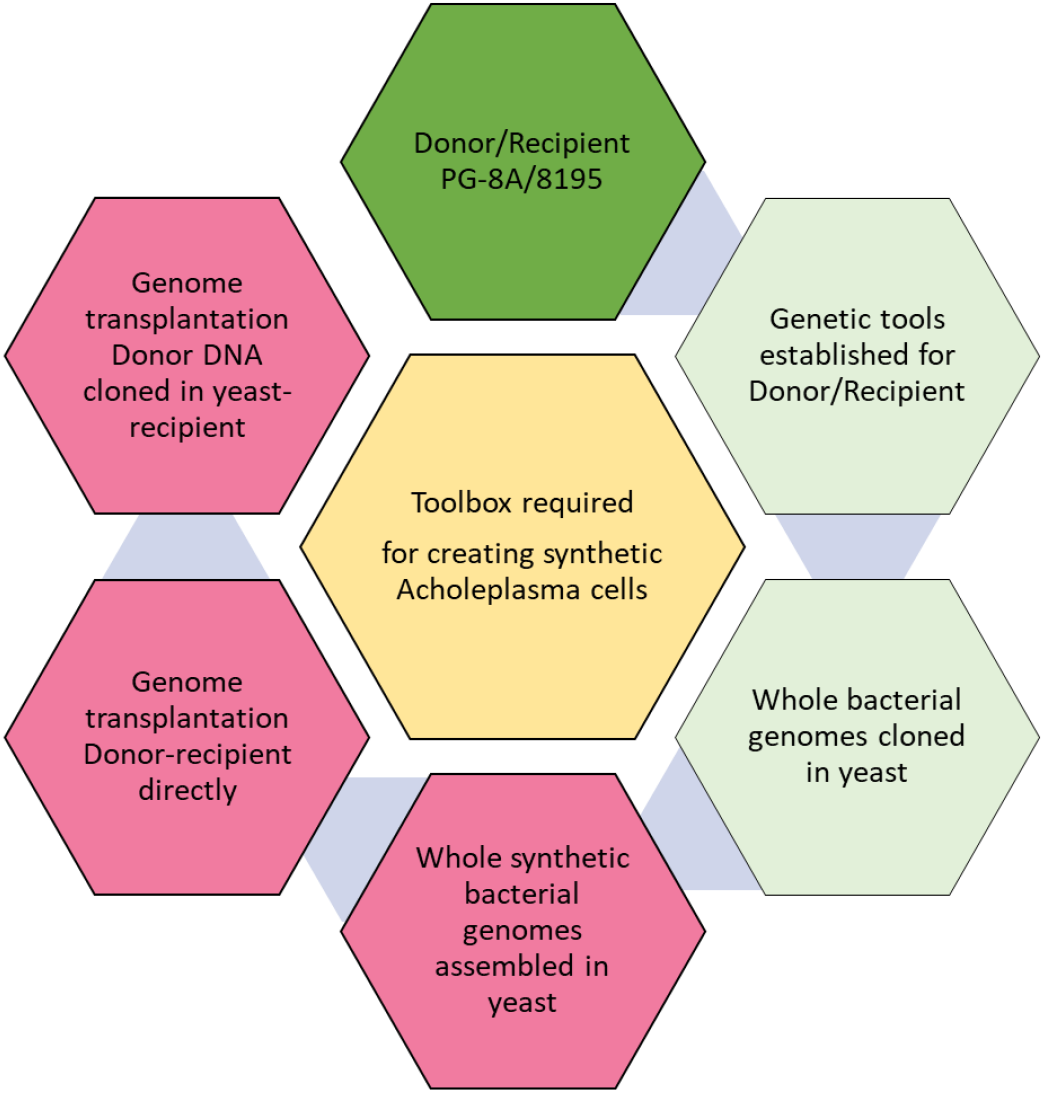
Genetic tools required for the creation of synthetic *Acholeplasma* strains. Dark green hexagon – proposed donor/recipient cells are selected. Light green hexagons – some tools exist. Red hexagons – all genetic tools for these tasks need to be developed.

First, we selected two *A. laidlawii* strains for use as a donor and recipient: PG-8A and 8195, respectively (Figure 2). Both strains, when grown without cholesterol, produce carotenoids, which result in the yellow color of pelleted cells (Figure 2A, B). Strain PG-8A has a darker pigmentation, which may serve as an initial visual screen following genome transplantation (Figure 2A, B). To gain a better understanding of the genetic differences between *A. laidlawii* PG-8A and 8195, the genome of each strain was sequenced using the Oxford Nanopore minION platform. The genomes of strains PG-8A and 8195 share 93% sequence identity, and the genomes are similar in size (1.5 Mbp) and have a G/C content of 32%. Furthermore, *A. laidlawii* strain 8195 has been described to lack restriction and recombination activities^18,23^, which makes it a good recipient strain. From our sequencing data, we identified components of a Type I restriction system in strain 8195. However, there is a nonsense mutation in the one restriction subunit identified, which would truncate the predicted protein at amino acid position 794/997. Interestingly, strain 8195 is derived from a parental *A. laidlawii* strain containing one active restriction system^18^. On the other hand, *A. laidlawii* PG-8A has five documented restriction systems (REBASE)^24^ that would need to be removed before this strain could be used as a recipient cell; therefore, we proposed to use it first as a donor.

**Figure 2.**
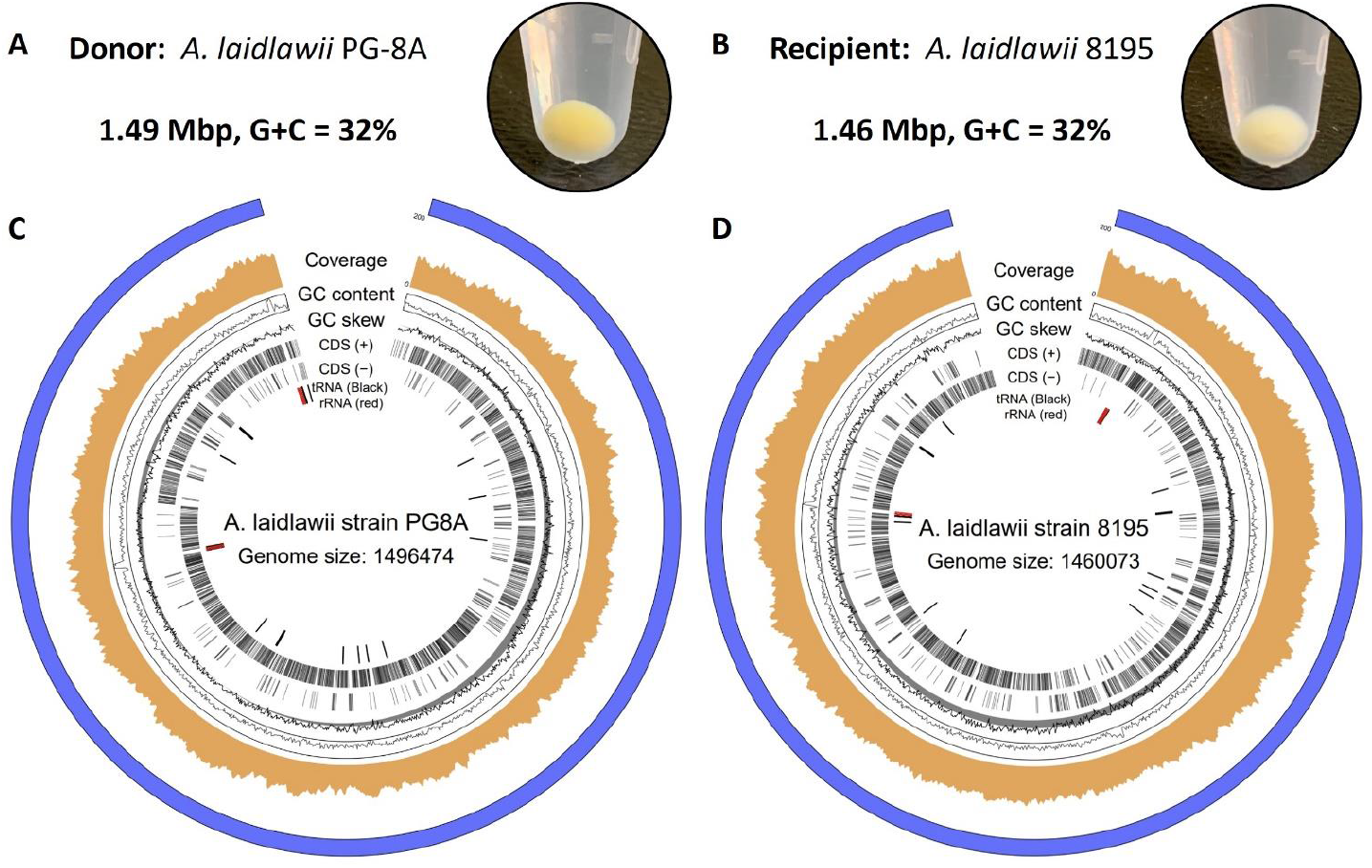
Proposed donor and recipient *Acholeplasma* strains for genome transplantation. Cell pellets and genome maps are shown for: A, C) *A. laidlawii* PG-8A (donor), B,D) *A. laidlawii* 8195 (recipient).

Furthermore, the PG-8A genome was already cloned in yeast^16^, which brings us one step closer to creating synthetic genomes in this host strain. In addition to restriction enzymes, many *Mollicutes* secrete nucleases to digest exogenous DNA^25–27^. The presence of potent extracellular nucleases may be an important consideration for future genome transplantation experiments, as the donor genome could be damaged prior to recipient-cell uptake. We developed an assay for easy evaluation of the activity of *A. laidlawii* nucleases (Figure 3), similar to what has been done with some *Mollicutes* previously^25^. This assay involved incubating live *A. laidlawii* cells with plasmid DNA, after which the nucleases were inactivated with ethylenediaminetetraacetic acid (EDTA)^27^ and the cell/DNA mix was visualized on an agarose gel. Indeed, we observed digestion of extracellular plasmid DNA when incubated with live *A. laidlawii* cells of either strain (Figure 3). Removal of these nucleases using targeted or random mutagenesis may be necessary to generate a suitable recipient strain.

**Figure 3.**
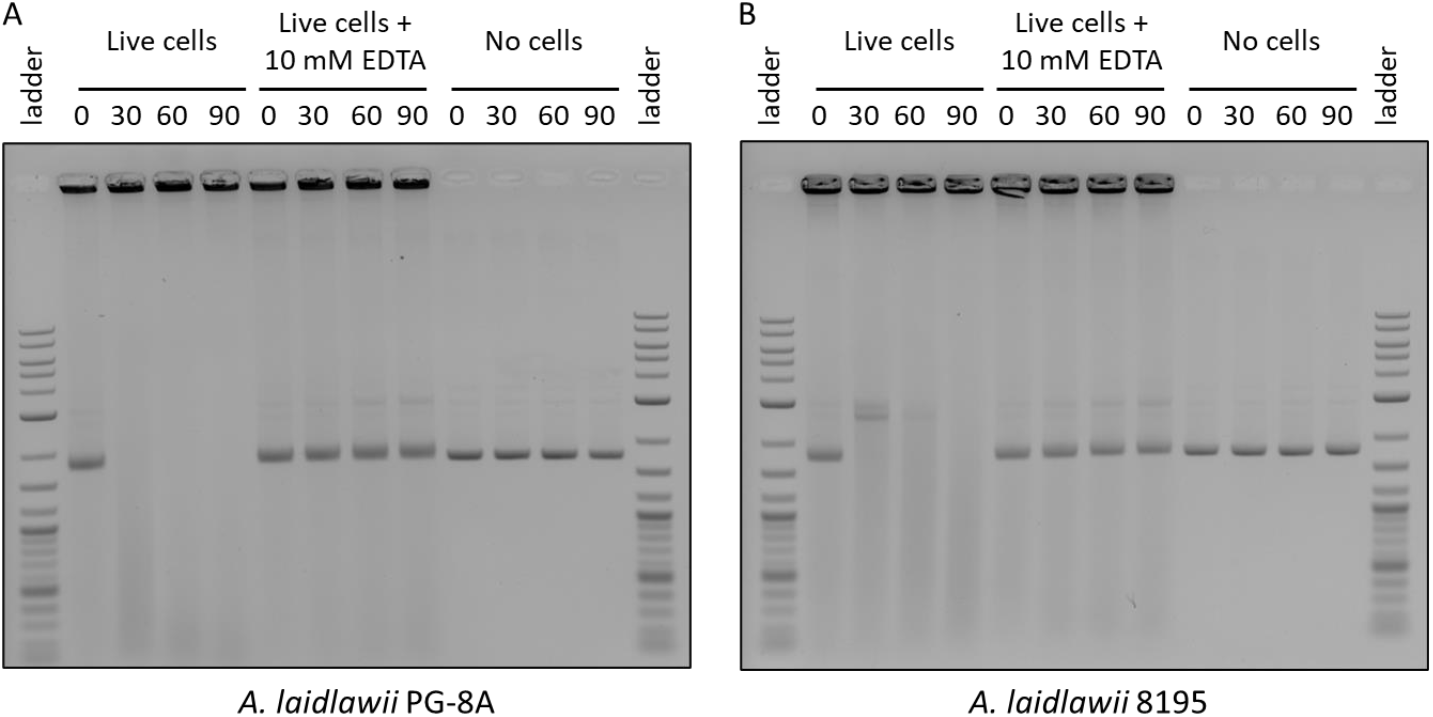
An assay demonstrates the potency of *A. laidlawii* extracellular nucleases. Live *A. laidlawii* cells were incubated with pUC19 DNA and incubated at 34°C for 0 – 90 minutes prior to inactivation with 10 mM ethylenediaminetetraacetic acid (EDTA). As a control, another set of samples was inactivated with EDTA before incubation. The cell/DNA mixes were visualized on an agarose gel. (A) Results of PG-8A assay. (B) Results of 8195 assay.

With the demonstration that EDTA can inhibit these nucleases, we set out to optimize an electroporation protocol. To this end, we first created a new shuttle plasmid that can replicate in yeast, *Escherichia coli*, and *A. laidlawii*. Artificial plasmids have been developed for many *Mollicute* species by cloning genomic regions containing DnaA boxes into vectors^28^. We followed the same strategy by cloning the putative origin of replication (OriC) of *A. laidlawii* 8195, which contains DnaA boxes upstream of the DnaA gene (Supplementary Figure 1), to create plasmid pAL1 (Figure 4A). For selection in *A. laidlawii*, we included a tetracycline (TetM) and puromycin resistance gene. In case the plasmid could not be replicated in *A. laidlawii*, we also included two 500-bp regions of homology to the 8195 genome flanking a homolog of ACL_0117 in strain PG-8A, which is partially toxic to yeast^16,29^.

**Figure 4.**
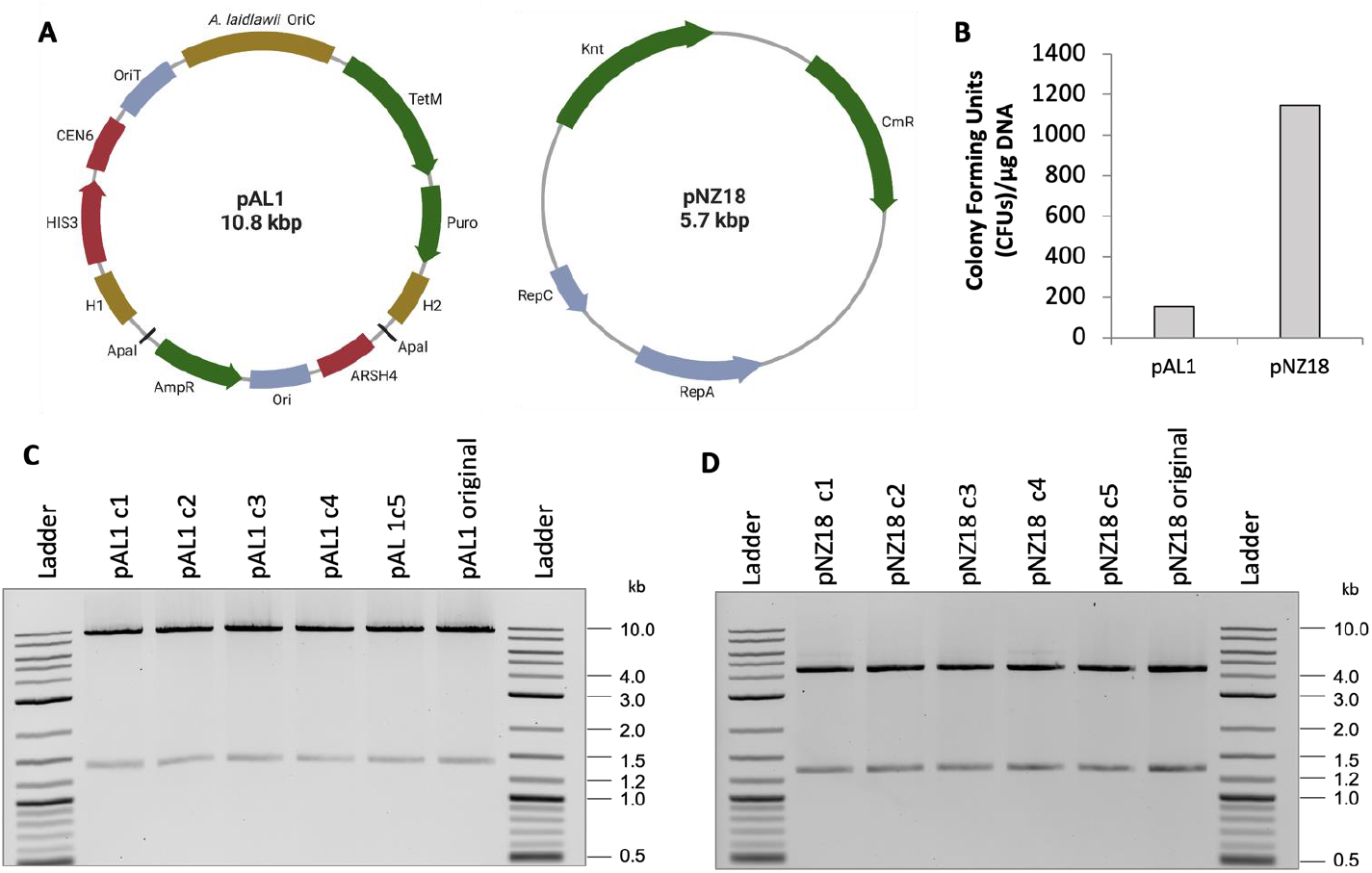
Overview of plasmid transformation to *A. laidlawii* 8195. A) Plasmid maps of pAL1 and pNZ18. OriC = Origin of replication, OriT = origin of transfer, AmpR = ampicillin resistance gene, Ori = pMB1 replicon, H1/H2 = regions of homology to *A. laidlawii* 8195 genome, Puro = puromycin resistance gene, TetM = tetracycline resistance gene, Knt = kanamycin/neomycin resistance gene, CmR = chloramphenicol resistance gene. Maps were created with BioRender.com. B) Colony counts of *A. laidlawii* 8195 following transformation with pAL1 or pNZ18. Bars represent an average colony count from 3 independent experiments. C) Restriction digests of pAL1 and pNZ18 plasmid DNA recovered from five *A. laidlawii* transformants. pAL1 original and pNZ18 original were the plasmids used for *A. laidlawii* transformation.

Next, we transferred pAL1 to *A. laidlawii* 8195 using previously reported PEG-mediated and electroporation protocols^30^. However, in our hands, these protocols were very inefficient and inconsistent. Subsequently, we optimized electroporation by changing volumes, temperatures, and the addition of yeast tRNA and EDTA (Supplementary Table 1), and we also considered what has been done in other protocols and with other *Mollicutes*^17,31,32^. Using the optimized electroporation protocol, we obtained transformation of pAL1 to *A. laidlawii* with a frequency of 2 × 10^−6^/colony forming unit (CFU) per 1 µg of DNA, or 1.56 × 10^2^ CFUs per 1 µg DNA. Transformation of pAL1 was only successful when selected for with tetracycline: the puromycin marker was not functional (data not shown). The puromycin marker uses a promoter from the Tuf gene of *Mycoplasma capricolum*, and it is possible that the promoter is weak or not recognized in *A. laidlawii*. For tetracycline selection, we used 1 µg/mL as this concentration allows for selection and still prevents the appearance of spontaneous mutations during the time when true transformants are selected (after 4 – 6 days). It is important to note that after 10 – 12 days, colonies started to appear on our negative control selection plates.

Following our success in transforming pAL1, we attempted our protocol with pNZ18^33^, a plasmid that contains a promiscuous gram-positive replicon (Figure 4A). This plasmid has previously been reported to replicate in *A. laidlawii*^21^. We obtained a transformation frequency of 3.7 × 10^−6^ CFU per 1 µg DNA, or 1.15 × 10^3^ CFUs per 1 µg DNA (Figure 4C), which is higher than what was initially reported for wildtype 8195^21^. We selected for transformants with 200 µg/mL neomycin to prevent the appearance of spontaneous mutants, which appeared within a similar timeframe to transformed *A. laidlawii* when selected at concentrations below 60 µg/mL. To enable assembly in yeast of constructs into pNZ18 for delivery to *A. laidlawii*, we added elements for selection and maintenance in *S. cerevisiae*. The new plasmid, called pNZ18-CAH, showed similar efficiency of electroporation compared to original pNZ18 (Supplementary Figure 2).

We recovered pAL1 and pNZ18 plasmids from *A. laidlawii* transformants grown in appropriate selective media by performing DNA isolation and transfer to *E. coli* to obtain a higher concentration of plasmids. Plasmids recovered from *E. coli* were digested, and we saw no gross rearrangements in 5/5 clones tested for each plasmid (Figures 4 C and D). Unlike pAL1, which was recovered in *E. coli* Epi300, pNZ18 was recovered in strain MC1061. Plasmids containing the pSH71 replicon, such as pNZ18, have been noted to have poor transformation efficiencies to RecA-minus *E. coli* strains^34^.

When *A. laidlawii* transformants were propagated in nonselective liquid media, we observed quick loss of both plasmids (Supplementary Table 2). This is a useful feature when propagation of a plasmid is required only for short period of time such as the delivery of genome-editing tools (example: Cas9). This experiment also indicated that pAL1 does not experience a high rate of genomic integration either within the OriC or at homology regions. This is consistent with previous unsuccessful attempts to integrate a plasmid into strain 8195, likely because this strain contains a premature stop codon in the RecA gene^23,35^.

Our transformation frequency of pNZ18 is an improvement to what has previously been reported by Sundström and Wieslander for wild type 8195^21^. However, following transformation and curing of pNZ18, this group had also reported the isolation of a strain that could be retransformed to a drastically higher frequency relative to wild type^21^. Curious if we could recreate such a strain with our own transformation protocol, we subjected wild type *A. laidlawii* 8195 to four consecutive rounds of electroporation and plasmid curing. We then isolated single colonies from the final pool of transformed cells and tested transformation frequency of several strains (Figure 5 A). Interestingly, we identified strains with noticeably improved electroporation frequency (Figure 5 B).Plasmids from evolved *A. laidlawii* could also be recovered in *E. coli* (data not shown).

**Figure 5.**
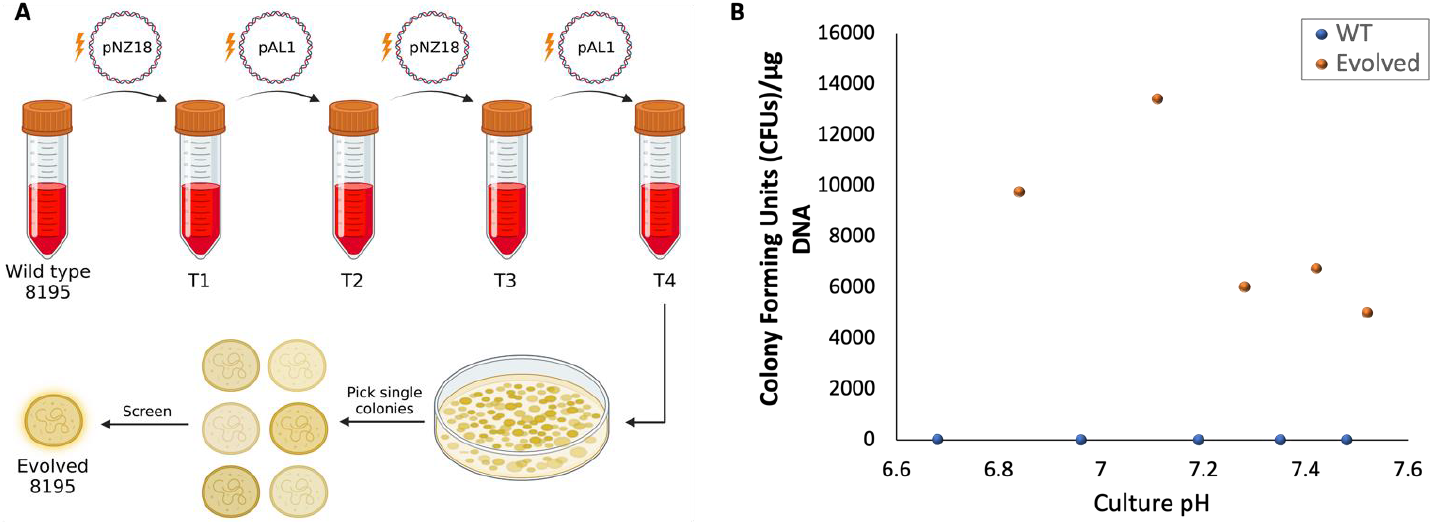
Creation of an *A. laidlawii* strain with improved electroporation frequency. A) Schematic of the evolution process. Following four rounds of electroporation and curing, single *A. laidlawii* colonies were screened for a higher transformation frequency. Created with BioRender.com B) Results of electroporation to wild type and a single evolved *A. laidlawii* 8195 strain. Both strains were transformed at various stages of growth, determined by pH of the culture, with 1 microgram of pAL1 DNA. Colony counts are from a single electroporation experiment.

Currently, PEG-mediated transformation is the only known way to facilitate genome transplantation. Although a method using PEG has previously been established for *A. laidlawii*^36^, the method differs from the traditional transplantation-like protocol, which involves relatively low concentrations of PEG and a longer incubation. We therefore aimed to adapt the transplantation-like protocol to *A. laidlawii* 8195. We first performed a series of pilot experiments to determine a starting protocol for further testing. Shown in Supplementary Table 4 is a summary of our PEG transformation experiments with cultures at various stages of growth and varied protocol parameters. Motivated by our previous success with evolution, many of these experiments were performed with a strain that had been isolated and cured from a previous PEG transformation. Overall, we got successful transformations in 27 out of 88 experiments. In future work, parameters from successful transformation will be used for future optimization of PEG transformation and transplantation protocols. Additionally, several more rounds of PEG transformation/curing could be performed to continue evolution of an improved recipient cell. Alternatively, a targeted approach, such as using a CRISPR base editor^37^, could be used to disrupt genes encoding putative extracellular nucleases or other genetic factors that could be affecting plasmid transformation/genome transplantation.

On the other hand, integration of genetic cassettes into the genome is an important tool on our road to create *A. laidlawii* driven by synthetic genomes. Therefore, we tested if Tn5 transposase can be used for this purpose. To this end, we first constructed a Tn5 cassette by PCR amplifying the TetM gene with primers that flanked the gene with 19-bp mosaic ends (Figure 6B). The cassette was mixed with EZ-Tn5 Transposase to generate a transposome, which was then electroporated to *A. laidlawii*. The protocol was the same except that we removed EDTA and yeast tRNA, which may negatively affect the transposase. We obtained transformants for both 8195 and PG-8A (Figure 6A), but in contrast to transformation with plasmid DNA, Tn5 transformations are less consistent (Supplementary Table 3). We have not noticed an increase in Tn5 transformation frequency when we use an evolved *A. laidlawii* strain. We created a second Tn5 cassette using the neomycin marker from pNZ18, but electroporation of this cassette into *A. laidlawii* has not produced any colonies. It is possible that the neomycin marker is not as effective when it is integrated into the genome as a single copy, but more transformations should be performed before this is concluded. Further experiments will be necessary to optimize transposon integration into the genome. We confirmed the presence of the Tn5-TetM transposon in PG-8A by passing the transformed strains multiple times to remove any original DNA and then performing PCR analysis (Figure 6C). In future work, we plan to use strain PG-8A transformed with Tn5 as a donor to first establish bacteria-to-bacteria genome transplantation followed by transplantation of a PG-8A genome that has been cloned in yeast. Furthermore, large-scale mutagenesis will be performed to create a derivative of strain 8195 with reduced or abolished nuclease activity for use as a recipient.

**Figure 6.**
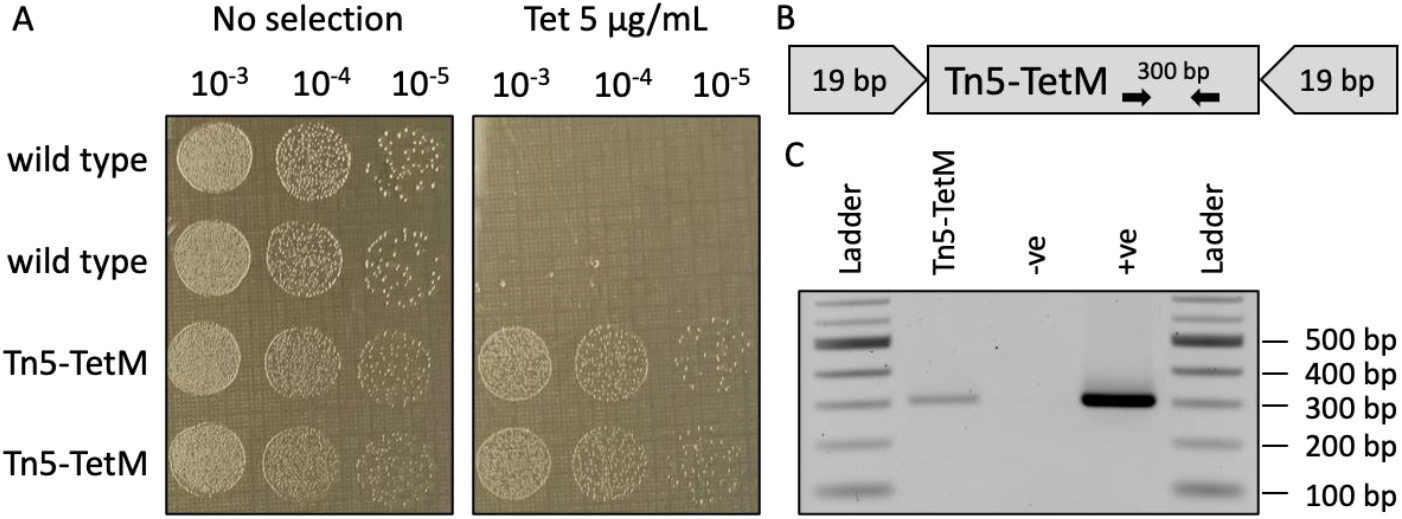
Tn5 transposase insertion into *A. laidlawii* PG-8A to generate donor strain. A) Serial dilutions of wildtype (WT) PG-8A and one colony transformed with Tn5 transposase (Tn5-TetM) were spot plated on SP-4 plates without selection or SP-4 containing 5 μg/mL tetracycline. B) Schematic of the Tn5 transposon cassette. The tetracycline resistance gene (TetM) is flanked by 19-bp mosaic ends that are recognized by the Tn5 transposase. PCR primers for colony screening that bind to the TetM gene are shown. C) Confirmation of marker insertion in donor *A. laidlawii*. PCR amplification with primers that bind to TetM produces a band of the expected size in Tn5-TetM *A. laidlawii* and the original Tn5 cassette (+ve), but not in the WT strain (-ve).

In conclusion, we have developed an expanded *A. laidlawii* genetic toolbox, including replicative plasmids, an improved transformation protocol, an improved strain for DNA uptake via electroporation, and a method of genomic integration, which will enable our future work towards the goal of creating synthetic/hybrid strains that should open new possibilities to study *Acholeplasmas* and possibly *Phytoplasmas*. In the meantime, our multi-host plasmid can be used to clone/assemble genetic pathways in yeast that can be then moved by electroporation to be propagated as episomes in *A. laidlawii* 8195.

## MATERIALS AND METHODS

### Strains and Cultures

*A. laidlawii* strain 8195 was kindly provided by Dr. Kevin Dyvbig and Dr. James Daubenspeck at the University of Alabama at Birmingham. *A. laidlawii* strains PG-8A (ATCC 23206) and 8195 were grown in SP-4 media lacking fetal bovine serum (3.5 g/L BBL mycoplasma broth base, 10 g/L tryptone, 5.3 g/L peptone, 8.8 g/L yeast extract, 5 g/L glucose, 0.6 g/L L-glutamine, 1.1 g/L NaHCO_3_, 2 g/L yestolate, 4 mL 0.5% phenol red solution, adjusted to pH = 7.6 with NaOH and sterilized through filtration) at 32°C or 34°C without shaking. For PEG transformation experiments, SP-4 was supplemented with 17% horse serum. Solid media was made with 0.95 - 1% agar and without phenol red solution. In addition to appropriate antibiotics (1 μg/mL tetracycline or 200 μg/mL neomycin), SP-4 was always supplemented with 200 u/mL penicillin.

*E. coli* strains Epi300 (Lucigen, Cat #: LGN-EC300110) and MC1061 (NCBI: txid 1211845) were grown in Luria Broth (LB) (5 g/L yeast extract, 10 g/L tryptone, 10 g/L NaCl) at 37°C, which was supplemented with 10 μg/mL tetracycline or 10 μg/mL chloramphenicol when appropriate. Solid LB was prepared with 1.5% agar.

*S. cerevisiae* VL6-48 (ATCC MYA 3666) was grown in 2x YPAD (100 g/L YPD broth, 160 mg/L adenine hemisulfate salt) at 30°C. Following transformation, *S. cerevisiae* was instead grown in a synthetic drop-out media lacking histidine containing 1 M sorbitol.

### Preparation of Tn5 Transposomes

The TetM gene, with its native promoter and terminator, was PCR-amplified from pAL1 using 5’-phosphorylated primers with 19-bp flanking mosaic ends (ME) that are recognized by the Tn5 transposase (Supplementary Table 5). The resulting PCR product was purified and concentrated using EZ-10 Spin Columns (BioBasic). The purified product was diluted to a final concentration of 100 ng/μL in TE buffer, and it was then combined with EZ-Tn5 transposase (Lucigen) as described in the manufacturer’s instructions. 1 μL of the prepared transposome was used for transformation to *A. laidlawii*.

### Transformation

#### Transformation to A. laidlawii

*A. laidlawii* culture was grown to OD_600_ = 0.2 - 0.25. Prior to harvesting, 1 mL aliquots of culture were pre-treated with 50 μL of 100 mM EDTA and incubated at 34°C for 10 minutes. Cells were then centrifuged at 15,596 x g for 10 minutes at room temperature. Supernatant was discarded, and cell pellets were resuspended in 1 mL of room-temperature wash buffer (272 mM sucrose, 8 mM HEPES, pH = 7.4). Cells were spun as before, and supernatant was removed. The pellet was resuspended in 100 μL wash buffer and kept on ice for 5 minutes. 5 μg of pAL1 DNA or 1 μg of pNZ18 DNA (dissolved in ddH_2_O) and 10 μg of yeast tRNA was added to cells, and the mix was incubated on ice for 2 - 3 minutes. The cell/DNA mix was then transferred to a pre-chilled 2 mm cuvette (VWR) and pulsed in a GenePulser Xcell (Bio-Rad) at 2.5 kV for 5 ms (200 Ω, 25 μF). Cells were recovered in 1 mL of ice-cold SP-4 and kept on ice for 10 minutes. The cells were then transferred to a 1.5 mL tube and incubated at 34°C for 2 hours before plating. Plates were kept at 34°C for 4-6 days. For the transformation of transposomes, EDTA pretreatment and yeast tRNA were omitted. For transformations comparing wild type and evolved *A. laidlawii*, cells were grown and recovered at 32°C, only 1 microgram of pAL1 DNA was used, and the use of yeast tRNA and EDTA was omitted.

#### Evolution of A. laidlawii

Wild type *A. laidlawii* 8195 was subjected to four rounds of electroporation, alternating between pNZ18 and pAL1. Electroporation was performed as described above, but cells were grown and recovered at 32°C, and the EDTA pretreatment step and the use of yeast tRNA were excluded. After each round, transformants were pooled in a liquid culture and passed several times without antibiotic selection to promote plasmid curing. Following the fourth round of transformation and passing, serial dilutions of the resulting culture were plated on non-selective SP-4 agar media to obtain single colonies. Colonies were screened for plasmid loss and were tested for improved transformation frequency against the wild type strain.

#### Nuclease Assay

Cultures of *A. laidlawii* PG-8A and 8195 were grown to OD_600_ = 0.2. Cultures were centrifuged at 15,596 x g for 10 minutes, and the cell pellet was concentrated in 1/10^th^ the original volume with wash buffer. 20 μL of concentrated cells were combined with 630 ng of pUC19 DNA in a final volume of approximately 40 μL. For the experimental condition, the cell/DNA mix was incubated at 34°C for 0, 30, 60, or 90 minutes. After the appropriate incubation time, EDTA was added to the mix at a final concentration of 10 mM to stop nuclease activity. For the EDTA control, the same amount of EDTA was instead added prior to incubation. For the DNA-only condition, 20 μL of wash buffer was used in place of cells, and EDTA was not added. Following all incubation times, each reaction was run on a 1% TAE agarose gel and imaged with ethidium bromide.

## ASSOCIATED CONTENT

### Supporting Information

The Supplementary File contains Supplementary Figure 1, Supplementary Tables 1 – 5, and Supplementary Methods.

## Author Contributions

B.J.K. and D.P.N. conceived the experiments, D.P.N., N.D.M., D.J.G., and B.J.K. conducted the experiments, D.P.N., N.D.M., D.J.G., and B.J.K. analyzed the results, D.P.N., and B.J.K. wrote the paper. All authors edited the paper. All authors have given approval to the final version of the manuscript.

## Notes

The authors declare no competing financial interests.

The genome sequence of *A. laidlawii* 8195 has been deposited to NCBI (BioProject PRJNA975591). Plasmids pAL1 and pNZ18-CAH are available on Addgene (ID 197285 and ID 203159, respectively).

## ACKNOWLEDGMENT

This work was supported by the Government of Canada’s New Frontiers in Research Fund (NFRF), [NFRFE-2018-01124]. In addition, research in B.J.K. laboratory is also supported by Natural Sciences and Engineering Research Council of Canada (NSERC), [RGPIN-2018-06172]. Plasmid pNZ18 and sequence/annotation data were kindly provided by the lab of Dr. Leendert Hamoen at the University of Amsterdam. The graphical abstract was created by Emma J. L. Walker.

