## Supplemental for "An expanded genetic toolbox to accelerate the creation of *Acholeplasma laidlawii* driven by synthetic genomes"

aatcaaaaaaccacctttcttatgaaaccttgcttttcttattataaataaagtgtacttaaaagtcaaacataaaatggtctttcttttatttatttttttatttgatttttcccacaatt
tttaatatgaatgttccccacaattattgtccacacattgtggataaagtttccacatttttattcacaatgttgataagtagcgtaagtatatttaacagccttacaaagcaaatga
tacactgaaagttatccacaatttaatatttaaagaacagctaaatcaaaaagttatccacaaataatgtggaaaactttttattaaatttgtcgtttcttatgctatcatagtttta
cataaattattaactcagggaggcagtcatgagtccaaacagcacactatggcagacaatattacaggatttagaaaaactatacaacgaggagacttacaacgagctatttc
taccagtgacttctacttttaaagatcaaaacggattacttacaatggttgtagcaaatgagttcttaaagaatcgtatcaataaactatacatcgcaaaaattaacgaacttgc
tactaaatattcaagtactccagttagattgaaattcatatcacaagaggaagttattgaagaaccagtagcggatcgtaaattaaccattgattatcgtcaaggtaacttaaac
tctacatatacctttgactcttttgttgttggaaaatctaacatgtttgcttttcgtatggcgatgaaggttgctgatcaacctgcagcagtagcaaacccccttctacatatttggtg
atgtaggtttaggtaaaacccatcttatgcaagcaataggtaactatatattagataatgatgttgaaaaacgtatattatatgttaaagctgataattttattgaagactttgtat
cattattatcaagaaacaaaaataagactgaagaattcaatgctaaatataaagatatagatgttatattagtggatgatattcaaattatggccaacgctagtaaaactcaaa
tggaattctttaaactctttgactatctatatttaaataataagcaaattgtcattacgtctgataaaccagcttcacaattaacaaatattatgccgcgattaacaacacgttttg
aagctggtctctctgtagacatacaaatacctgaattagaacatagaataagcattttaaagagaaaaacagctacattagatgcaaacttagaggtaggagaggatatctta
acctttattgcatctcaatttgcagcaaatattagagaaatggagggtgcactcattcgtttaattagttatgcacaaacctttaatctagaaataacaatggatgttgttgaaga
agcacttggtgctgtcttaaaaacaaagaagaaaacaaatcaattaaacgaaaataactacgataagatccaaagtatcgttgcagattacttccaagtgtcattaccagactt
aattggtaaaaaaagacatgctaaattcacattacctagacatatcgcaatgtatcttatcaaactcaaattcaatataccttataaaacgattggttctttgtttaatgatagag
accactctactgtattggctgcttgtgagaaagtagaacgcgatatgaggatggattcgaacttaaagtttgctgttgactcaattgtcaaaaaaatagattcaccatcattaaa
gtgataaatgtttataaaaatgattaaatgtggtaaaataaatggtagatgaagcgattattgctagtttcccactttcccacagacactaacaataacaaagaagaaataata
attaataaaggggtaaaat

**Supplementary Figure 1. Sequence of the *A. laidlawii* 8195 origin of replication.** The CDS of the DnaA gene is shown in grey, with the start codon highlighted in green and the stop codon highlighted in red. Highlighted in yellow are DnaA boxes predicted by Ori-Finder 2022.

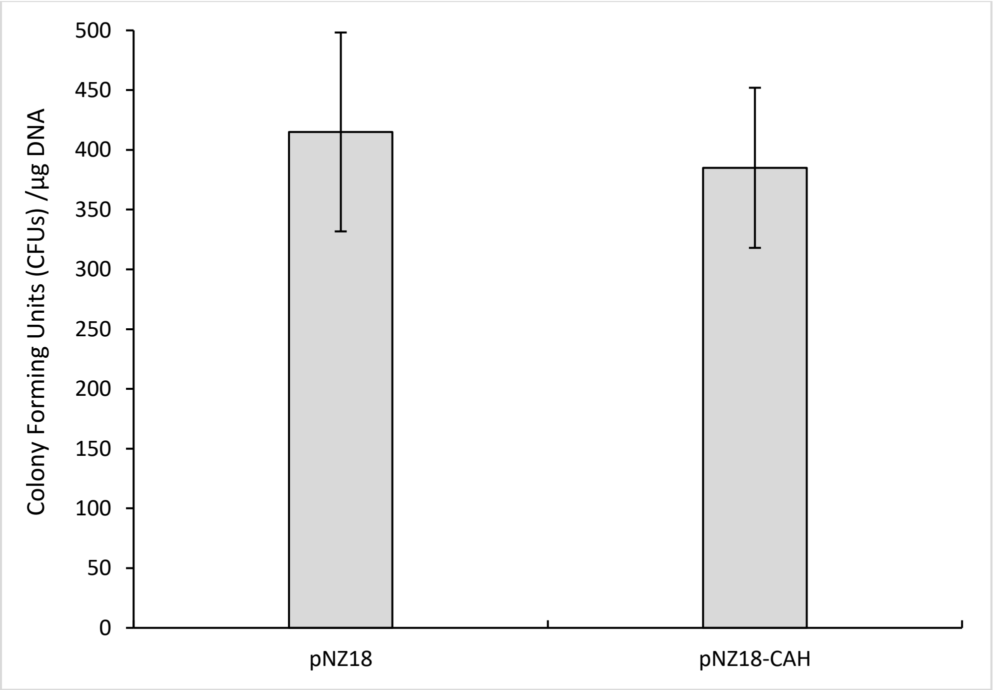


**Supplementary Figure 2.** Comparison between pNZ18 and pNZ18-CAH. Bars represent the average colony counts from 4 independent electroporation experiments to an evolved *A. laidlawii* 8195 strain using 1 microgram of pNZ18 or pNZ18-CAH, with error bars showing standard error of the mean. The difference in mean colony counts between the two plasmids is not statistically significant, as calculated by a two-tailed student's t-test.

**Supplementary Table 1. Optimization of *A. laidlawii* transformation protocol**. Each section represents an independent transformation of pAL1 to strain 8195. All experiments were done with cultures of $OD_{600}$ = 0.17 - 0.25, with a recovery time of at least 2 hours at 34°C following transformation. All colonies were counted after 6 days at 34°C.

| Condition Tested | Colony Count |
|---|---|
| **Volume of Culture Used (mL)** | |
| 1 | 44 |
| 1.5 | 8 |
| 2 | 7 |
| 3 | 6 |
| **Number of Washes Before Transformation** | |
| 1 | 194 |
| 2 | 10 |
| 3 | 11 |
| **Amount of DNA Transformed (µg) and use of 10 µg yeast tRNA (+ or -)** | |
| 2 - | 11 |
| 2 + | 2 |
| 3 - | 20 |
| 3 + | 30 |
| 5 - | 34 |
| 5 + | 36 |
| 10 - | 45 |
| 10 + | 90 |
| **Cell Pretreatment and Incubation (1)** | |
| Pretreatment- 5 µg/mL chloramphenicol for 1 hour | 5 |
| Pretreatment- 5 µg/mL tetracycline for 1 hour | 4 |
| Pretreatment- 0.25% trypsin for 10 mins | 0 |
| Pretreatment- ~1 mM EDTA for 10 mins | 92 |
| Pretreatment- ~2 mM EDTA for 10 mins | 111 |
| Pretreatment- ~5 mM EDTA for 10 mins | 107 |
| Pretreatment- 30 mins on ice | 360 |
| Electroporation- 400 Ω instead of 200 Ω | 70 |
| After electroporation- 10 mins on ice | 266 |
| After electroporation- 10 mins at room temperature | 62 |
| **Cell Pretreatment and Incubation (2)** | |
| After electroporation- 10 mins at room temperature | 220 |
| After electroporation- 10 mins on ice | 329 |
| Pretreatment- ~5 mM EDTA for 10 mins | 306 |
| Pretreatment: ~5mM EDTA for 10 mins + After transformation: 10 mins on ice | 432 |

**Supplementary Table 2. Stability of pAL1 and pNZ18 in *A. laidlawii.*** Plasmid stability was analyzed after 50 hours of growth at 37°C.

| pNZ18 | | | | | |
|---|---|---|---|---|---|
| Grown with antibiotic | | | Grown without antibiotic | | |
| No selection | Neo 200 | Ratio | No selection | Neo 200 | Ratio |
| 158 | 180 | 114 | 613 | 60 | 10 |
| **pAL1** | | | | | |
| Grown with antibiotic | | | Grown without antibiotic | | |
| No selection | Tet 1 | Ratio | No selection | Tet 1 | Ratio |
| 184 | 132 | 72 | 725 | 70 | 10 |

**Supplementary Table 3. Summary of Tn5 transformation experiments to *A. laidlawii*.**

| Experiment # | Strain | $OD_{600}$ | Recovery time (hours) | Antibiotic Selection (µg/mL) | # of Colonies |
|---|---|---|---|---|---|
| 1 | 8195, WT | 0.2699 | 4 | Tet1 | 220 |
| 2 | 8195, WT | 0.2000 | 2 | Tet1 | 11 |
| 3 | 8195, WT | 0.0900 | 2 | Tet1 | 0 |
| 4 | 8195, WT | 0.2297 | 2 | Tet1 | 0 |
| 5 | 8195, WT | 0.2077 | 4 | Tet1 | 0 |
| 6 | 8195, WT | 0.2216 | 4 | Tet0.75 | 17 |
| 7 | PG-8A | 0.2200 | 2.67 | Tet1 | 1 |
| 8 | 8195, Evolved | ~0.25 | 2 | Tet1 | 23 |

**Supplementary Table 4**. Summary of PEG transformation experiments to *A. laidlawii*. Shown for each experiment is the strain used, growth of cultures as determined by pH, the volume of cells plated after the transformation, the plasmid used for each transformation, and the colony counts for each technical replicate. Along with each group of experiment is the list of parameters changed from the standard protocol described in the methods section. Strains: WT = wild type 8195, E = strain evolved for improved electroporation, PEG = a strain transformed and cured from an earlier PEG experiment. SP4- : SP-4 lacking serum; SP4+: SP-4 supplemented with 17% horse serum. N/D – no data

| # | Date | Strain | Culture pH | Volumed Plated (µL) | Plasmid | Rep #1 | Rep #2 | Rep #3 | Protocol Notes |
|---|---|---|---|---|---|---|---|---|---|
| 1 | 2023-04-26 | PEG | 7.27 | 500 | pAL1 | 0 | N/D | N/D | • 3-hour PEG incubation<br>• 2-hour recovery |
| 2 | 2023-04-26 | PEG | 7.37 | 500 | pAL1 | 0 | N/D | N/D | |
| 3 | 2023-04-26 | PEG | 7.32 | 500 | pAL1 | 0 | N/D | N/D | |

| | | | | | | | | | |
|---|---|---|---|---|---|---|---|---|---|
| 4 | 2023-04-26 | E | 7.19 | 500 | pAL1 | 0 | N/D | N/D | |
| 5 | 2023-04-26 | E | 7.26 | 500 | pAL1 | 0 | N/D | N/D | |
| 6 | 2023-04-26 | E | 7.38 | 500 | pAL1 | 0 | N/D | N/D | |
| 7 | 2023-05-04 | PEG | 7.20 | 500 | pAL1 | 6 | N/D | N/D | • Cultures were diluted from SP4+ to SP4+<br>• 1.5-hour PEG incubation<br>• 2-hour incubation |
| 8 | 2023-05-04 | PEG | 7.35 | 500 | pAL1 | 10 | N/D | N/D | |
| 9 | 2023-05-04 | PEG | 7.43 | 500 | pAL1 | 20 | N/D | N/D | |
| 10 | 2023-05-04 | OC2 | 7.12 | 500 | pAL1 | 1 | N/D | N/D | |
| 11 | 2023-05-04 | E | 7.32 | 500 | pAL1 | 0 | N/D | N/D | |
| 12 | 2023-05-04 | E | 7.40 | 500 | pAL1 | 0 | N/D | N/D | |
| 13 | 2023-05-12 | PEG | 7.53 | 500 | pAL1 | 20 | N/D | N/D | • Cultures were diluted from SP4+ to SP4+<br>• 1.5-hour PEG incubation<br>• 2-hour recovery |
| 14 | 2023-05-12 | PEG | 7.65 | 500 | pAL1 | 125 | N/D | N/D | |
| 15 | 2023-05-12 | PEG | 7.69 | 500 | pAL1 | 35 | N/D | N/D | |
| 16 | 2023-05-12 | PEG | 7.44 | 500 | pAL1 | 36 | N/D | N/D | • Cultures were diluted from SP4+ to SP4+<br>• 1.5-hour PEG incubation<br>• 2-hour recovery<br>• Used a pool of cured PEG-transformed strains |
| 17 | 2023-05-12 | PEG | 7.61 | 500 | pAL1 | 5 | N/D | N/D | |
| 18 | 2023-05-12 | PEG | 7.57 | 500 | pAL1 | 0 | N/D | N/D | |
| 19 | 2023-05-19 | WT | 6.89 | 500 | pAL1 | 0 | N/D | N/D | • Cultures were diluted from SP4- to SP4+<br>• 2-hour PEG incubation<br>• 1.5-hour recovery |
| 20 | 2023-05-19 | WT | 7.14 | 500 | pAL1 | 2 | N/D | N/D | |
| 21 | 2023-05-19 | WT | 7.29 | 500 | pAL1 | 0 | N/D | N/D | |
| 22 | 2023-05-19 | WT | 7.41 | 500 | pAL1 | 0 | N/D | N/D | |
| 23 | 2023-05-19 | WT | 7.53 | 500 | pAL1 | 0 | N/D | N/D | |

| | | | | | | | | | |
|---|---|---|---|---|---|---|---|---|---|
| 24 | 2023-05-19 | E | 6.69 | 500 | pAL1 | 2 | N/D | N/D | |
| 25 | 2023-05-19 | E | 7.05 | 500 | pAL1 | 14 | N/D | N/D | |
| 26 | 2023-05-19 | E | 7.27 | 500 | pAL1 | 0 | N/D | N/D | |
| 27 | 2023-05-19 | E | 7.43 | 500 | pAL1 | 2 | N/D | N/D | |
| 28 | 2023-05-19 | E | 7.54 | 500 | pAL1 | 0 | N/D | N/D | |
| 29 | 2023-05-19 | WT | 6.89 | 500 | pAL1 | 0 | N/D | N/D | • Cultures were diluted from SP4- to SP4+<br>• 1.5-hour PEG incubation<br>• 2-hour recovery |
| 30 | 2023-05-19 | WT | 7.14 | 500 | pAL1 | 0 | N/D | N/D | |
| 31 | 2023-05-19 | WT | 7.29 | 500 | pAL1 | 0 | N/D | N/D | |
| 32 | 2023-05-19 | WT | 7.41 | 500 | pAL1 | 0 | N/D | N/D | |
| 33 | 2023-05-19 | WT | 7.53 | 500 | pAL1 | 0 | N/D | N/D | |
| 34 | 2023-05-19 | WT | 6.89 | 500 | pNZ18 | 0 | N/D | N/D | |
| 35 | 2023-05-19 | WT | 7.14 | 500 | pNZ18 | 0 | N/D | N/D | |
| 36 | 2023-05-19 | WT | 7.29 | 500 | pNZ18 | 0 | N/D | N/D | |
| 37 | 2023-05-19 | WT | 7.41 | 500 | pNZ18 | 0 | N/D | N/D | |
| 38 | 2023-05-19 | WT | 7.53 | 500 | pNZ18 | 0 | N/D | N/D | |
| 39 | 2023-05-19 | E | 6.69 | 500 | pAL1 | 0 | N/D | N/D | |
| 40 | 2023-05-19 | E | 7.05 | 500 | pAL1 | 1 | N/D | N/D | |
| 41 | 2023-05-19 | E | 7.27 | 500 | pAL1 | 1 | N/D | N/D | |
| 42 | 2023-05-19 | E | 7.43 | 500 | pAL1 | 1 | N/D | N/D | |
| 43 | 2023-05-19 | E | 7.54 | 500 | pAL1 | 0 | N/D | N/D | |
| 44 | 2023-05-19 | E | 6.69 | 500 | pNZ18 | 1 | N/D | N/D | |

| | | | | | | | | | |
|---|---|---|---|---|---|---|---|---|---|
| 45 | 2023-05-19 | E | 7.05 | 500 | pNZ18 | 0 | N/D | N/D | |
| 46 | 2023-05-19 | E | 7.27 | 500 | pNZ18 | 0 | N/D | N/D | |
| 47 | 2023-05-19 | E | 7.43 | 500 | pNZ18 | 0 | N/D | N/D | |
| 48 | 2023-05-19 | E | 7.54 | 500 | pNZ18 | 0 | N/D | N/D | |
| 49 | 2023-06-01 | WT | 7.29 | 200 | pAL1 | 0 | 0 | N/D | • Cultures were diluted from SP4+ to SP4+<br>• 3-hour PEG incubation<br>• 1-hour recovery |
| 50 | 2023-06-01 | WT | 7.38 | 200 | pAL1 | 0 | 0 | N/D | |
| 51 | 2023-06-01 | WT | 7.49 | 200 | pAL1 | 0 | 0 | N/D | |
| 52 | 2023-06-01 | WT | 7.64 | 200 | pAL1 | 0 | 0 | N/D | |
| 53 | 2023-06-01 | E | 7.23 | 200 | pAL1 | 0 | 0 | N/D | |
| 54 | 2023-06-01 | E | 7.32 | 200 | pAL1 | 0 | 0 | N/D | |
| 55 | 2023-06-01 | E | 7.40 | 200 | pAL1 | 0 | 0 | N/D | |
| 56 | 2023-06-01 | E | 7.54 | 200 | pAL1 | 0 | 0 | N/D | |
| 57 | 2023-06-01 | PEG | 7.43 | 200 | pAL1 | 0 | 0 | N/D | |
| 58 | 2023-06-01 | PEG | 7.48 | 200 | pAL1 | 0 | 0 | N/D | |
| 59 | 2023-06-01 | PEG | 7.62 | 200 | pAL1 | 0 | 0 | N/D | |
| 60 | 2023-06-01 | PEG | 7.74 | 200 | pAL1 | 0 | 0 | N/D | |
| 61 | 2023-06-06 | PEG | 7.42 | 200 | pAL1 | 0 | N/D | N/D | • Culture was diluted from SP- to SP+<br>• 2x Fusion Buffer contained 250 mM NaCl<br>• 2-hour PEG incubation<br>• 1.5-hour recovery |

| | | | | | | | | | |
|---|---|---|---|---|---|---|---|---|---|
| 62 | 2023-06-06 | PEG | 7.42 | 200 | pAL1 | 1 | N/D | N/D | • Culture was diluted from SP- to SP+<br>• 2x Fusion Buffer contained 250 mM NaCl<br>• 2-hour total PEG incubation: 1-hour incubation without DNA, 1-hour incubation after DNA was added<br>• 1.5-hour recovery |
| 63 | 2023-06-06 | PEG | 7.42 | 200 | pAL1 | 0 | N/D | N/D | • Culture was diluted from SP- to SP+<br>• 2-hour PEG incubation<br>• 1.5-hour recovery |
| 64 | 2023-06-06 | PEG | 7.42 | 200 | pAL1 | 8 | N/D | N/D | • Culture was diluted from SP- to SP+<br>• 2-hour total PEG incubation: 1-hour incubation without DNA, 1-hour incubation after DNA was added<br>• 1.5-hour recovery |
| 65 | 2023-06-07 | WT | 7.13 | 200 | pAL1 | 0 | 0 | 0 | • Cultures were diluted from SP- to SP+<br>• 2-hour PEG incubation<br>• 1.5-hour recovery |
| 66 | 2023-06-07 | WT | 7.27 | 200 | pAL1 | 0 | 0 | 0 | |
| 67 | 2023-06-07 | WT | 7.47 | 200 | pAL1 | 0 | 0 | 0 | |
| 68 | 2023-06-07 | E | 7.31 | 200 | pAL1 | 0 | 0 | 0 | |
| 69 | 2023-06-07 | E | 7.50 | 200 | pAL1 | 0 | 0 | 0 | |
| 70 | 2023-06-07 | E | 7.65 | 200 | pAL1 | 0 | 0 | 0 | |
| 71 | 2023-06-07 | PEG | 7.10 | 200 | pAL1 | 0 | 0 | 0 | |
| 72 | 2023-06-07 | PEG | 7.29 | 200 | pAL1 | 0 | 0 | 0 | |
| 73 | 2023-06-07 | PEG | 7.46 | 200 | pAL1 | 0 | 0 | 0 | |
| 74 | 2023-06-20 | PEG | 6.90 | 200 | pAL1 | 0 | 0 | 0 | • Cultures were diluted from SP- to SP+ |

| | | | | | | | | | |
|---|---|---|---|---|---|---|---|---|---|
| 75 | 2023-06-20 | PEG | 7.18 | 200 | pAL1 | 0 | 0 | 0 | • 3-hour total PEG incubation: 1-hour incubation without DNA, 2-hour incubation after DNA was added<br>• 1-hour recovery |
| 76 | 2023-06-20 | PEG | 7.35 | 200 | pAL1 | 0 | 0 | 0 | |
| 77 | 2023-06-20 | PEG | 7.47 | 200 | pAL1 | 1 | 2 | 1 | |
| 78 | 2023-06-20 | PEG | 7.58 | 200 | pAL1 | 0 | 0 | 0 | |
| 79 | 2023-06-21 | PEG | 6.93 | 200 | pAL1 | 0 | 0 | 0 | • Cultures were diluted from SP- to SP+<br>• 2-hour total PEG incubation: 1-hour incubation without DNA, 1-hour incubation after DNA was added<br>• 1.5-hour recovery |
| 80 | 2023-06-21 | PEG | 7.15 | 200 | pAL1 | 0 | 0 | 0 | |
| 81 | 2023-06-21 | PEG | 7.34 | 200 | pAL1 | 1 | 0 | 0 | |
| 82 | 2023-06-21 | PEG | 7.46 | 200 | pAL1 | 1 | 1 | 0 | |
| 83 | 2023-06-21 | PEG | 7.55 | 200 | pAL1 | 5 | 1 | 1 | |
| 84 | 2023-06-23 | PEG | 7.01 | 200 | pAL1 | 0 | 0 | 0 | • Cultures were diluted from SP- to SP+<br>•2-hour total PEG incubation: 1-hour incubation without DNA, 1-hour incubation after DNA was added<br>• 1.5-hour recovery |
| 85 | 2023-06-23 | PEG | 7.19 | 200 | pAL1 | 1 | 0 | 0 | |
| 86 | 2023-06-23 | PEG | 7.36 | 200 | pAL1 | 1 | 0 | 0 | |
| 87 | 2023-06-23 | PEG | 7.44 | 200 | pAL1 | 2 | 1 | 1 | |
| 88 | 2023-06-23 | PEG | 7.52 | 200 | pAL1 | 2 | 0 | 0 | |

**Supplementary Table 5. List of primers used for the creation and screening of the Tn5 TetM cassette.** Underlined sequences show 19-bp mosaic ends.

| Primer Name | Sequence (5' - 3') | Description |
|---|---|---|
| BK2348_F | ctgtctcttatacacatctgcggccgctccctttagtgagggttaatgtcgt | Primer for creating Tn5 cassette |
| BK2348_R | ctgtctcttatacacatctgctactagtattattatgtgattttgttgaa | Primer for creating Tn5 cassette |

| Primer Name | Sequence (5' - 3') | Size (bp) | Description |
|---|---|---|---|
| BK2580_F | tcacgcattacgtaaaatgg | 300 bp | Primer for screening TetM gene |
| BK2580_R | aaagaacagttttggaaacgaa | 300 bp | Primer for screening TetM gene |

**Supplementary Table 6. List of primers used in the construction of pAL1.** Underlined sequences show parts of the primer that bind to the template, where the remaining sequence is an overhang that adds an overlapping region of homology.

| Name | Sequence (5' - 3') | Description |
|---|---|---|
| BK1970_F | acgtatcagtaaagatttaaaaagattaggatttaaatttgggcccgatcgccaacaaat | Amplifies ARSH4 |
| BK1970_R | caattaatgtgagttagctcactcattaggcaccccaggcggatcgcttgcctgtaactt | Amplifies ARSH4 |
| BK1971_F | aaaagatacgaggcgcgtgtaagttacaggcaagcgatccgcctggggtgcctaatgagt | Amplifies pMB1 replicon, AmpR |
| BK1971_R | taaatacaactttaccggttagcttaaccttatctccatcgggcccgcggaacccctatt | Amplifies pBM1 replicon, AmpR |
| BK1972_F | tgtatttagaaaaataaacaaataggggttccgcgggcccgatggagataaggttaagct | Amplifies Homology 1 |
| BK1972_R | caagctcttactatattaccctgttatccctagcgtaactcttaaaagaaagatactact | Amplifies Homology 1 |
| BK1973_F | ttcttttaagagttacgctagggataacagggtaatatagtaagagcttggtgagcgcta | Amplifies HIS3, CEN6 |
| BK1978_R | cgctataatgaccccgaagcagggttatgcagcggaagatttacatcttcggaaaacaa | Amplifies HIS3, CEN6 |
| BK1981_F | ataacaaagaagaaataataattaataaaggggtaaaattccctttagtgagggttaat | Amplifies TetM |
| BK1974_R | tcgacctgcaggcatgcaagcttggcgtaatcatggtcatgctactagtattattatgtg | Amplifies TetM |
| BK1975_F | cgttatatgttcaacaaaatcacataataatactagtagcatgaccatgattacgccaag | Amplifies LacZα |
| BK1975_R | tatatattttacacttatttaatacttaattcaaaaaatactatgcggcatcagagcaga | Amplifies LacZα |
| BK1976_F | atggtgcactctcagtacaatctgctctgatgccgcatagtatttttgaattaagtatt | Amplifies Puro |
| BK1976_R | ctagttaaagtagacatctacatactccggaaaatctatatttaataataaaaaatcggg | Amplifies Puro |
| BK1977_F | taacaaaaaaatcgggaaatcccgatttttattattaaatatagattttccggagtatg | Amplifies Homology 2 |
| BK1977_R | agagcaaggtaaaaggtagtatttgttggcgatcgggcccaaatttaaatcctaatcttt | Amplifies Homology 2 |
| BK1979_F | aattaaagaaaaaatagtttttgttttccgaagatgtaaaatcttccgctgcataaccct | Amplifies OriT |
| BK1979_R | aagaaaagcaaggtttcataagaaaggtggtttttgattgatcgtcttgccttgctcgt | Amplifies OriT |
| BK1980_F | gagctgtaagtacatcaccgacgagcaaggcaagacgatcaatcaaaaaaccacctttct | Amplifies OriC |
| BK1980_R | agggttttcccagtcacgacattaaccctcactaaagggaattttaccccttttattaat | Amplifies OriC |

### Supplementary Methods

#### Whole-Genome Sequencing.

Cultures of *A. laidlawii* PG-8A and 8195 were grown to a high density, and genomic DNA was extracted using the Monarch High Molecular Weight DNA Extraction Kit (New England Biolabs). Genomic DNA was sequenced using a 9.4.1 Flongle flow cell with the RBK004 rapid barcoding kit following library preparation as instructed by the manufacturer (Oxford Nanopore Technologies). The Flongle flow cell was refueled during the sequencing run to improve output. Runs were basecalled using Guppy v4.5.2 in high-accuracy mode, and the resulting output was assembled using Flye v2.8.1. Genome maps were generated using a custom R script following annotation with Prokka and alignment with minimap2, samtools, and mosdepth[1–5]. Putative DnaA boxes were identified with Ori-Finder 2022[6]. For identifying restriction genes, draft genomes were annotated with RAST[7].

#### Screening for the Tetracycline Marker.

Wildtype and Tn5-TetM *A. laidlawii* were screened using Multiplex PCR (Qiagen) with primers listed in Supplementary Table 4. 1 µL of *A. laidlawii* genomic DNA or 1 ng of the Tn5 cassette was used as template. The thermocycler was programmed for 95°C for 5 minutes, 20 cycles of 94°C for 30 seconds, 57°C for 90 seconds, and 72°C for 90 seconds, followed by a final extension of 72°C for 10 minutes.

#### Plasmid Construction.

The plasmid pAL1 was constructed through homologous recombination of overlapping fragments in yeast. All plasmid parts were first PCR-amplified using PrimeSTAR GXL DNA Polymerase (Takara Bio) with primers that added 40 bp of overlapping homology to each end: assembly primers are listed in Supplementary Table 5. The fragments were then combined in roughly equimolar amounts and co-transformed into *S. cerevisiae* spheroplasts; preparation of spheroplasts has been described previously[8]. Following spheroplast recovery, plasmid DNA was isolated and transformed to *E. coli* Epi300. Individual *E. coli* colonies were screened for the correctly assembled plasmid. The high copy pMB1 replicon, ampicillin resistance gene, and lacZα sequences were amplified from pUC19. The CEN6 and HIS3 sequences were amplified from pDMI2.0 (Karas lab), and the ARSH4 sequence was amplified from pAGE1.0[9]. The tetracycline marker, TetM, was isolated from genomic DNA of *Mesoplasma florum* L1-T-ΔRE[10]. *A. laidlawii* sequences (homology 1 and 2, origin of replication) were amplified from *A. laidlawii* 8195 genomic DNA. The origin of transfer was amplified from Pmod-yeast oriT (Karas lab). The puromycin marker was amplified from a previous *A. laidlawii* plasmid version, which was originally amplified from Pmod-yeast (Karas lab).

#### *A. laidlawii* Plasmid Recovery and Digest.

Following transformation experiments, 20 *A. laidlawii* transformants from each plasmid were passed on solid media with the appropriate antibiotic. 10 colonies were then inoculated to 10 mL of SP-4 media. Once grown, cells were pelleted, and DNA was isolated using alkaline lysis with Buffers P1, P2, and P3 (Qiagen) followed by alcohol precipitation. Approximately 1 µL of isolated DNA (~20 – 200 ng) was used to transform *E. coli* as described previously: pAL1 was transformed to strain Epi300, and pNZ18 was transformed to strain MC1061. For each transformation, plasmid DNA was miniprepped from two *E. coli* colonies using EZ-10 Spin Columns (BioBasic). pAL1 and pNZ18 were then digested in a 20-µL reaction with 6 units of EcoRV-HF and 3 units of PshAI (New England Biolabs), respectively, at 37°C for approximately 1 hour.

#### Plasmid Stability Assay.

A single colony of *A. laidlawii* 8195 harboring either pAL1 or pNZ18 was grown in SP-4 supplemented with tetracycline and neomycin, respectively. Cultures were grown to a high density before being diluted to $OD_{600}$ = 0.01 in i) SP-4 supplemented with the appropriate selective antibiotic and ii) SP-4 without selective antibiotic. After 50 hours of growth at 37°C, cultures were serially diluted and plated on solid SP-4 with and without selective antibiotics. All colonies were counted after 6 days at 37°C, and the ratio of colonies grown on non-selective and selective media was calculated.

**Transformation to *E. coli*.**

Electrocompetent *E. coli* were prepared by washing cells from 50 mL cultures ($OD_{600}$ = 0.5 - 0.8) three times with ice-cold $ddH_2O$ and concentrating cells in 10% glycerol. Typically, 30 µL of electrocompetent cells was combined with plasmid DNA in a 1.5 mL tube before transferring the cell/DNA mix to a pre-chilled 2 mm electrocuvette. The cuvette was pulsed using the same conditions described for *A. laidlawii*. Cells were recovered in 750 µL - 1 mL SOC media (5 mL 2M $MgCl_2$, 10 mL 250 mM KCl, 20 mL 1 M glucose, 0.5 g/L NaCl, 5 g/L yeast extract, 20 g/L Tryptone) with shaking at 225 rpm for 1 hour at 37°C before plating. For some transformations, electrocompetent *E. coli* Transformax Epi300 cells (Lucigen) were used.

**PEG Transformation to *A. laidlawii*.**

A culture of *A. laidlawii* 8195 was diluted in SP-4 supplemented with 17% horse serum and grown at 32°C for 16 – 24 hours. 1 mL of culture was centrifuged at 15,000 x g for 5 - 10 minutes at room temperature. Cells were washed with 1 mL of S/T Buffer (250 mM NaCl, 10 mM Tris-HCl [pH = 6.5]) and centrifuged as before. Supernatant was discarded; the cells were resuspended in 200 µL 0.1 M $CaCl_2$ and incubated on ice for 30 minutes. Next, the cells were added to a 50 mL conical tube containing 400 µL of SP-4 minus serum, 1 µL of yeast tRNA (10 µg/µL), 1 µg of plasmid DNA, and an equal volume of a modified 2x fusion buffer (500 mM NaCl, 20 mM $MgCl_2$, 20% PEG 8000, 20 mM Tris-HCl [pH = 6.5]: pH adjusted to 6.5). The cells were pipetted gently to mix and incubated at 32°C for 1 - 3 hours. 5 mL of prewarmed SP-4 plus serum was added, and the cells were left to recover at 32°C for 1 - 2 hours. Finally, the cells were centrifuged at 3,000 x g and 4°C for 20 minutes. Supernatant was discarded, the pellets were resuspended in 1 mL of SP-4 minus serum, and 200 - 500 µL was plated on SP-4 plates supplemented with appropriate antibiotics. Typically, colonies were counted after 6 days at 32°C.
